# Learning single-cell chromatin accessibility profiles using meta-analytic marker genes

**DOI:** 10.1101/2021.04.01.438068

**Authors:** Risa Karakida Kawaguchi, Ziqi Tang, Stephan Fischer, Chandana Rajesh, Rohit Tripathy, Peter K. Koo, Jesse Gillis

## Abstract

**Motivation:** Single-cell Assay for Transposase Accessible Chromatin using sequencing (scATAC-seq) is a valuable resource to learn cis-regulatory elements such as cell-type specific enhancers and transcription factor binding sites. However, cell-type identification of scATAC-seq data is known to be challenging due to the heterogeneity derived from different protocols and the high dropout rate.

**Results:** In this study, we perform a systematic comparison of 7 scATAC-seq datasets of mouse brain to benchmark the efficacy of neuronal cell-type annotation from gene sets. We find that redundant marker genes give a dramatic improvement for a sparse scATAC-seq annotation across the data collected from different studies. Interestingly, simple aggregation of such marker genes achieves performance comparable or higher than that of machine-learning classifiers, suggesting its potential for downstream applications. Based on our results, we reannotated all scATAC-seq data for detailed cell types using robust marker genes. Their meta scATAC-seq profiles are publicly available at https://gillisweb.cshl.edu/Meta_scATAC. Furthermore, we trained a deep neural network to predict chromatin accessibility from only DNA sequence and identified key motifs enriched for each neuronal subtype. Those predicted profiles are visualized together in our database as a valuable resource to explore cell-type specific epigenetic regulation in a sequence-dependent and -independent manner.

**Supplementary information:** Supplementary data are available at *xxxxxx* online.

**Key points:**

- 7 scATAC-seq datasets of mouse brain are systematically compared to benchmark the efficacy of neuronal cell-type annotation from gene sets.
- Redundant marker genes give a dramatic improvement for a sparse scATAC-seq annotation beyond the heterogeneity of scATAC-seq data.
- We reannotated all scATAC-seq data for detailed cell types using robust marker genes and their meta scATAC-seq profiles are publicly available at a new Meta scATAC-seq server.
- Predicted profiles from only DNA sequence using a deep neural network are visualized together to explore sequence-dependent and -independent epigenetic regulation.

## 1 Introduction

The elaborate developmental process of multicellular organisms relies on epigenetic marks encoded in a cell-type specific manner. Chromatin accessibility is an epigenetic signal that can be read out via high-throughput sequencing methods, reflecting the existence of active regulatory regions such as enhancers and promoters. Assay for Transposase-Accessible
Chromatin using sequencing (ATAC-seq) is a primary method to detect the epigenetic footprint of chromatin location through the insertion of barcode sequences by the Tn5 transposase [2]. Due to its high throughput and applicability, massive reference atlases of ATAC-seq have been constructed for diverse targets such as immune and neuronal cells [3–5]. However, bulk ATAC-seq analysis often measures the chromatin accessibility across a mixture of cell types, unless each cell-type is isolated in advance. Indeed, previous studies that analyzed ATAC-seq data for neuronal celltype specific epigenetic profiles used cell lines or recombinase driver lines [6], or micro-dissection of a specific region [7, 8]. Such approaches not only require laborious work to obtain each cell or driver line but also restrict the scope of the analysis to a biased set of cell types. In addition, independent processing and sequencing of each sample inevitably cause batch effects that may make it challenging to make comparisons across datasets.

In principle, single-cell ATAC-seq (scATAC-seq) can resolve many of the issues that are intrinsic to bulk ATAC-seq by obtaining the ATAC-seq profiles from a broader sampling of individual cells. One promising application of scATAC-seq is the identification of cell-type specific gene regulation performed by multiple proximal and distant enhancers [9–12]. To assign each chromatin accessibility profile to the transcriptome space of known cell types, it is common practice to estimate the gene activity profile by summing the read counts around the transcription start site (TSS) and/or entire gene body for each gene. However, in general, gene activity inferred from epigenetic profiles cannot be perfectly aligned to RNA-seq because assessing epigenetic profiles at the gene body alone ignores the complex regulation producing transcripts [4]. Moreover, the limited number of chromosome copies (e.g., 2 for diploid organisms) results in observed signals that are nearly binary (0, 1, or 2) and are sparsely located across genomic loci, making it difficult to obtain robust results from single datasets [13]. Integration of data to improve robustness is one solution to this noise and sparsity problem, but there is still no gold standard approach that grapples with the profound differences of scATAC-seq datasets across protocols [14]. To fully reveal cell-type specific gene regulation from scATAC-seq data, we need to develop methods to group shared cell-types and then uncover the key regulatory features that are robust.

Despite the comparatively unbiased characterization of most single cell methods, marker-based annotation has retained a strong role in the validation of cell clusters as real “types”. However, marker-based annotation is not immune to “dropout” in general, which frequently arises especially for single-cell analyses due to technical artifacts or stochastic expression of mRNA transcripts. A potential solution to address the two main shortcomings of scRNA-seq data, dropout and batch effect, is to analyze the marker genes that are co-expressed, in addition to distinctly expressed. This approach expands the repertoire of markers, making it unlikely that all of them would be dropped out in unison, thereby transforming a hard classification problem into a soft classification problem [15]. In essence, scRNA-seq experiments vastly expand our capacity to discover marker sets and it is plausible this can be used to build a model applicable to scATAC-seq data, where pure clustering is otherwise more challenging (involving weak genome-wide trends).

In this study, we carried out a comprehensive benchmark of celltype classification for mouse brain scATAC-seq datasets based on five marker gene sets. The mouse brain is one of the most complex systems and also one of the most heavily assayed, providing a useful test bed for assessment. We collected marker sets from previous scRNA-seq and scATAC-seq studies, as well as meta-analytic marker sets inferred from multiple scRNA-seq datasets. In a broad evaluation of marker sets, learning methods, and datasets, we aim to simply characterize which factors drive characterization performance. Our principal finding is that careful selection of marker genes, especially when they are chosen to increase redundancy meta-analytically, can greatly improve performance; this occurs to such a degree that if an adequately strong marker set is selected, simple aggregation of the gene-specific scATAC-seq signal characterizes cell-type remarkably well. This finding provides an important basis for future data integration and downstream applications of scATAC-seq analysis. Moreover, we trained a deep convolutional neural network (CNN) to classify which cell types are accessible for an input DNA using the largest scATAC-seq dataset collected in this study. Those predictions are published at our server with all collected and re-annotated datasets as well as marker sets for the integrative analysis with future scATAC-seq datasets.

## 2 Results

### 2.1 Benchmarking the influence of marker gene selection on cell-type resolution across scATAC-seq datasets

Figure 1A shows a general scATAC-seq analysis workflow including the differences between experimental protocols and quantification methods. While each experimental technology relies on a different principle, the common downstream analysis is that the cells are gathered into clusters according to the similarity of the chromatin accessibility for each bin or region around TSS and gene body of each gene, called gene activity. To infer the cell-type specific regulatory network from scATAC-seq profiles, it is essential to assign each chromatin accessibility profile to a cell type, either through known marker genes or by mapping to cell types inferred from transcriptome data. Averaging of gene activity profiles for each cluster, which is a common step for cluster-level cell typing, reduces the influence of stochastic noise, decreases the sparsity of the dataset, and requires fewer computational resources, when compared to the single-cell analysis. Then cluster-level cell typing is done by checking the cluster-specific enrichment of biomarker genes based on prior knowledge of existing cell types as shown in Figure 1B. When a cell type is inferred for each cluster, however, the resolution of cell typing is limited by the cluster size. This especially matters for brain scATAC-seq analyses, which potentially contain hundreds of cell types, and clusters are expected to contain several finer grained cell types.

**Fig.1.**
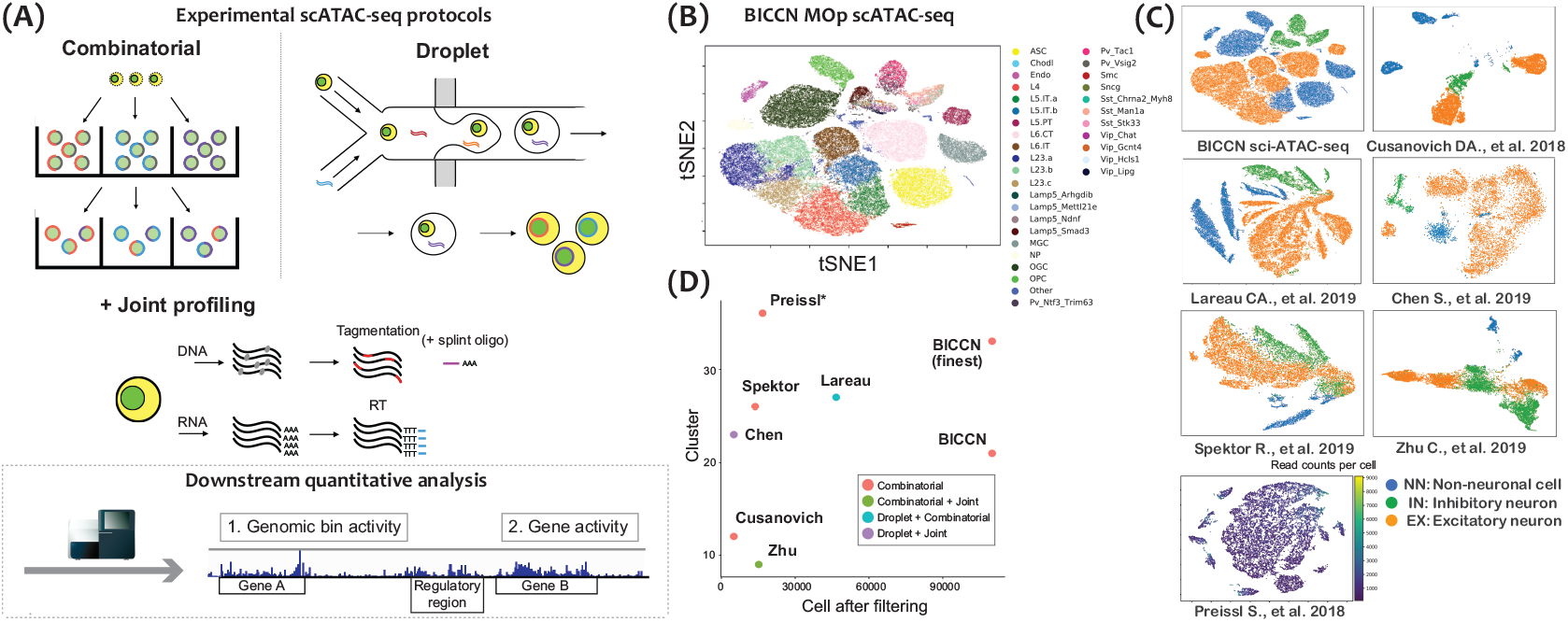
A workflow of scATAC-seq analysis and the seven scATAC-seq dataset used in this study. (A) A general workflow of scATAC-seq protocols and quantification. (B) tSNE mapping of BICCN scATAC-seq data from mouse primary motor cortex (MOp). (C) tSNE mapping of 7 scATAC-seq datasets used in this study. Each cell is colored depending on the assigned cell type from three major cell types; blue for non-neuronal cell (NN), green for inhibitory neuron (IN), and orange for excitatory neuron (EX). For the dataset without a cell-type annotation from [1], the total read count of each cell is shown by a different color. (D) Relationship between the number of detected clusters and sampled cells after filtering based on the coverage for 7 scATAC-seq datasets.

In fact, the disparity in terms of the number of cells and clusters for the heterogeneous datasets used in this study suggests that cluster-level annotation is inadequate since it would likely depend on an implausible lack of variation between pipelines. For example, Figure 1C and D shows the variation in their granularity (9 to 36 clusters), from major cell types to more detailed cell types, such as Pvalb or Vip within inhibitory neuronal cells. While characterization of individual cells, or at most small cell groups, is necessary for functional analysis, this strategy is only feasible with high-coverage datasets, such as the BICCN dataset, which has more than 110,000 cells. Besides, there is variation in marker gene selection, and thus definition of brain cell types, limiting annotation to major cell types, such as inhibitory (IN), excitatory (EX), and non-neuronal (NN) cell types,

To examine the real feasibility of our cell typing approach, therefore, we take two steps; benchmarking major cell-type annotation and its robustness at the cluster and individual cell level, then validating the annotation of more detailed cell types that are shared in two high-coverage datasets (Fig. 2A). Because the appropriate parameters or data handling processes for each study are unknown, we used the annotations provided by the authors to define “true” cell-type labels. For six out of seven datasets, the metadata about these cell types are published, or were provided personally. Among the seven datasets, three datasets are coupled with scRNA-seq data, one from the BICCN dataset [16] and two joint profiling data from Chen et al. [17] and Zhu et al. [18]. Those scRNA-seq data are later applied to validate the cell typing with the reference.

**Fig.2.**
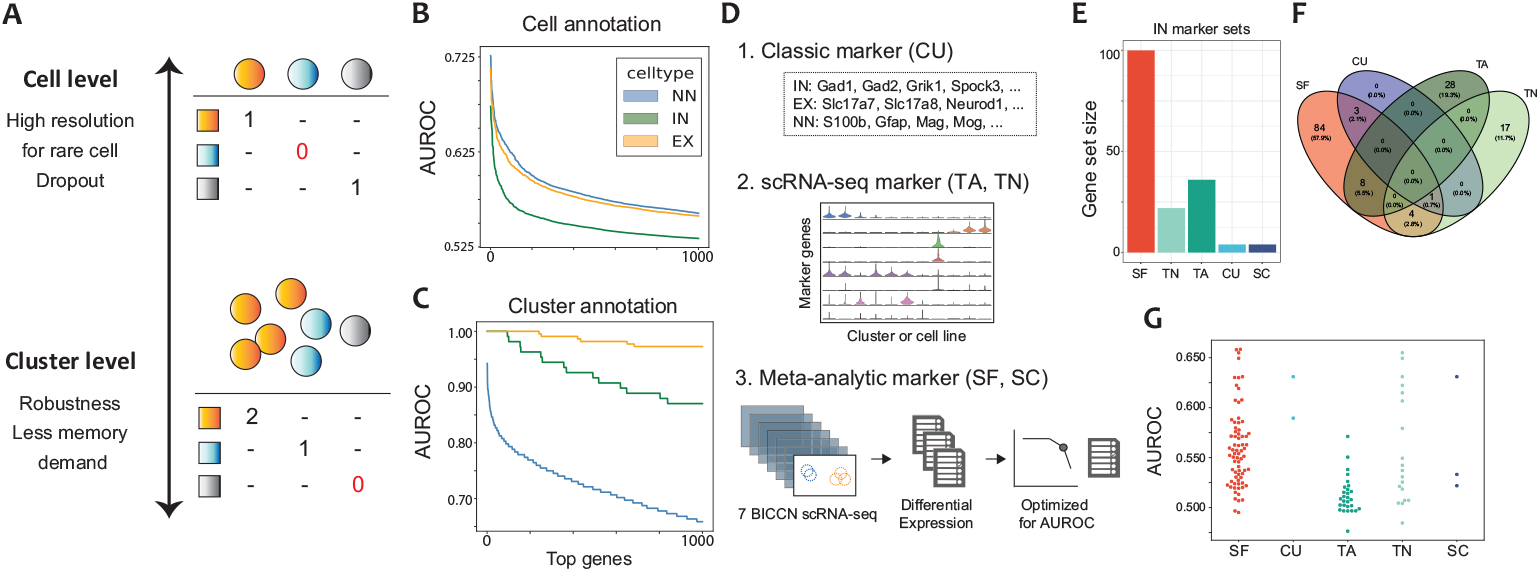
Cell-type characterization workflow at the cluster and individual cell level. (A) Graphical abstract of the cell-type annotation for scATAC-seq data using a single marker gene for each cell type. A dropout is observed with equal probability at cell level annotation while the cell types of the large cell cluster are more likely to be detected than rare cell types at cluster level. (B) and (C) The top 1,000 features for major cell-type classification for the BICCN scATAC-seq dataset using the gene activity of each single gene are shown for each cell (B) and cluster (C). (D) Classification of five marker sets used in this study. (E) and (F) The size and overlap of 5 marker gene lists for the IN cell type used in this study. (G) The distribution of AUROCs of the IN cell-type classification within the BICCN dataset using the gene activity of 5 marker gene sets.

### 2.2 Gene-set based approach using functional marker genes has a strong potential to produce practical and reproducible predictions at single-cell level

Due to the sparseness and binary-like characteristics, scATAC-seq data is prone to stochastic loss. To assess cell-type classification performance of each gene beyond such noises, area under the receiver operating characteristics curves (AUROCs) are computed for gene activity in the BICCN scATAC-seq dataset among the marjor categories: IN, EX, and NN cell types. The informativeness of individual genes, on average, only provides a modest predictive power of cell-types from random as expected (AUROC 0.5-0.6, Supplementary Fig. 4A). For example, by focusing on the top 1,000 most predictive genes, we find that fewer than 200 genes achieved an AUROC greater than 0.625 for each cell-type (Fig. 2B). On the other hand, aggregating gene activity at a cluster-level substantially improves the AUROC (Fig. 2C). It should be noted cell-level and cluster-level classification have different sample sizes (number of cells vs number of clusters), leading to a step-wise characteristic for cluster-level performance.

Additionally, we computed the AUROC for cell typing based on each genomic bin activity at cell-level classification (Supplementary Fig. 4B), as the features used in the previous studies [19]. Evidently, the activity level specified by top bins has the lowest predictive power with an AUROC < 0.55. We also computed p-values for Fisher’s exact test and around 6.39% of bins had p-values smaller than 0.05 after Bonferroni correction for three major cell types (inset in Supplementary Fig. 4B). These results show that no single feature provides sufficient accuracy, even for major cell types.

Previously, we found that using an expanded marker set that includes co-expressed genes of marker genes significantly improves cell typing in terms of signal-to-noise ratio and AUROCs for scRNA-seq data [15]. Thus, we hypothesize that this meta-analytic marker gene set (defined by scRNA-seq data) could also help improve the performance of cell typing with scATAC-seq data. To examine the validity of this idea, we collected five marker gene sets established for single-cell sequencing data, named *SF, CU, TA, TN*, and *SC*, for meta-analytic integration of brain scATAC-seq data (Fig. 2D-F). Specifically, the biomarker set determined in [12] (referred to as CU) is used as a scATAC-seq oriented biomarker set while those from [20] and [6] (TA and TN) are selected as the representatives of scRNA-seq oriented biomarker sets. Additionally, the gene sets SF and SC are newly constructed using six scRNA-seq datasets obtained in a BICCN project detailed in [16]. In Figure 2G, the AUROCs of marker genes computed for IN cell-types are substantially higher than random (0.5), particularly for our meta-analytic marker set SF. These distributions are not sufficiently high for practical cell-typing, but noticeably different from those of all genes, suggesting the potential predictive power of individual marker genes.

An extra advantage of the gene-set based approach is that the integrated signals are expected to be more robust for the difference between the omics layers. For example, the gene activity measured by scATACA-seq is known to imperfectly correspond to transcriptome profiles because it ignores complex regulation by distant or condition-specific enhancers [4]. However, the comparison of the gene activity profiles between the BICCN scRNA-seq and scATAC-seq as well as proteome data [21] shows that the overlaps of top cluster-specific genes can designate the cluster pairs of the consistent cell types better than the correlation coefficients for all captured genes (Supplementary Fig. 3A and D).

Taken together, our results indicate that a small number of marker genes substantially and consistently increase their activity in a specific cell type while most genes do not – though no single gene provides sufficient predictive power for consistent cell typing at individual cell level.

### 2.3 A meta-analytic marker gene set shows consistently increased signal across heterogeneous scATAC-seq datasets

We further evaluated the reproducibility of the AUROCs of single features across the different scATAC-seq datasets. By computing the Spearman’s correlation of AUROC scores for all pairs of datasets, we found large variation in the correlation between datasets (Supplementary Fig. 5). For example, the BICCN, *Lareau*, and *Chen* datasets were highly correlated with each other, while the *Cusanovich* and *Zhu* datasets had a much lower correlation with any dataset. Compared to the cell-level classification, the correlation coefficients of cluster-level or bin-level classification performances are comparable or even lower (e.g., comparisons involving the BICCN dataset), suggesting that the enrichment of high-performance features at cluster-level is only weakly reproducible across the datasets.

Focusing on the performance stability of the marker genes, we also computed the correlation for each marker gene set between the BICCN and *Lareau* datasets, which give us the highest correlation among all pairs of datasets for gene-level comparison. To evaluate statistical significance, we generated a null distribution by randomly sampling gene sets of the same size from all genes 10,000 times, and computed the correlation coefficients for each between the two datasets. As a result, all marker sets for IN type showed a higher correlation coefficient compared to the comparison of all genes (SF: 0.931, CU: 0.921, TA: 0.779, TN: 0.851, and SC: 0.880), but only the p-value of SF is significantly lower after multiple corrections (p-value<5e-5, *n* = 5). Together, these results demonstrate that cell typing by a single feature, such as gene activity or genomic bin, is highly variable at both individual cell and cluster-level across scATAC-seq datasets. At the same time, choosing meta-analytic marker genes can greatly increase the reproducibility of cell-type classification in scATAC-seq data.

### 2.4 Redundant and meta-analytic marker gene sets enable robust and practical cell-type classification

Following our evaluation of individual genes, we now consider the integration of information across multiple genes. To address this problem in a practical scATAC-seq analysis workflow, we evaluated the performance of cluster-level annotation based on two cell-typing strategies; the qualitative comparison of cluster-specific genes, and quantitative comparison of the aggregated pseudo-bulk profiles of gene activity.

As a cell-type classification using a gene list, the Jaccard index was computed as a metric of overlap between each marker set and cluster-specific genes for all clusters from seven datasets. Specifically, the index is defined as 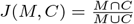, where *M* and *C* are the marker gene set and cluster-specific genes, respectively. Then, the indices of each cluster were normalized within the range (0, 1) across three cell types as shown in Figure 3A. Finally, we examined AUROCs based on the normalized Jaccard index for each cell type classification against the reference true celltype labels for the clusters. In Figure 3B, the AUROCs of each marker set are shown as a function of the number of top cluster-specific genes selected (shown in the x-axis). The larger the number of cluster-specific genes is, the higher the AUROCs of prediction are for all marker sets. However, the classification based on the SF marker set shows a sudden increase of the AUROCs to 0.8-0.9 only with around the top 100 cluster-specific genes. Descent performance of SC compared to SF indicates the importance of marker set redundancy for scATAC-seq annotation. Interestingly, most of the marker sets except for TA reached around 0.8 for the classification of IN cell type just by considering the top genes. This result suggests that even a few genes are enough to accurately annotate the IN cell-type group while additional genes can further improve the accuracy of prediction for other cell types. As a redundant gene set which is comparable to SF, we compared the AUROCs with those based on the cell-type specific gene sets inferred from each dataset (Supplementary Fig. 7). The SF marker set produced highest and most stable AUROCs across the different cell types compared to most of the gene sets, suggesting the potential of our meta-analytic approach to enhance the stability and applicability of marker sets beyond the cell-type and batch-specific difference.

**Fig.3.**
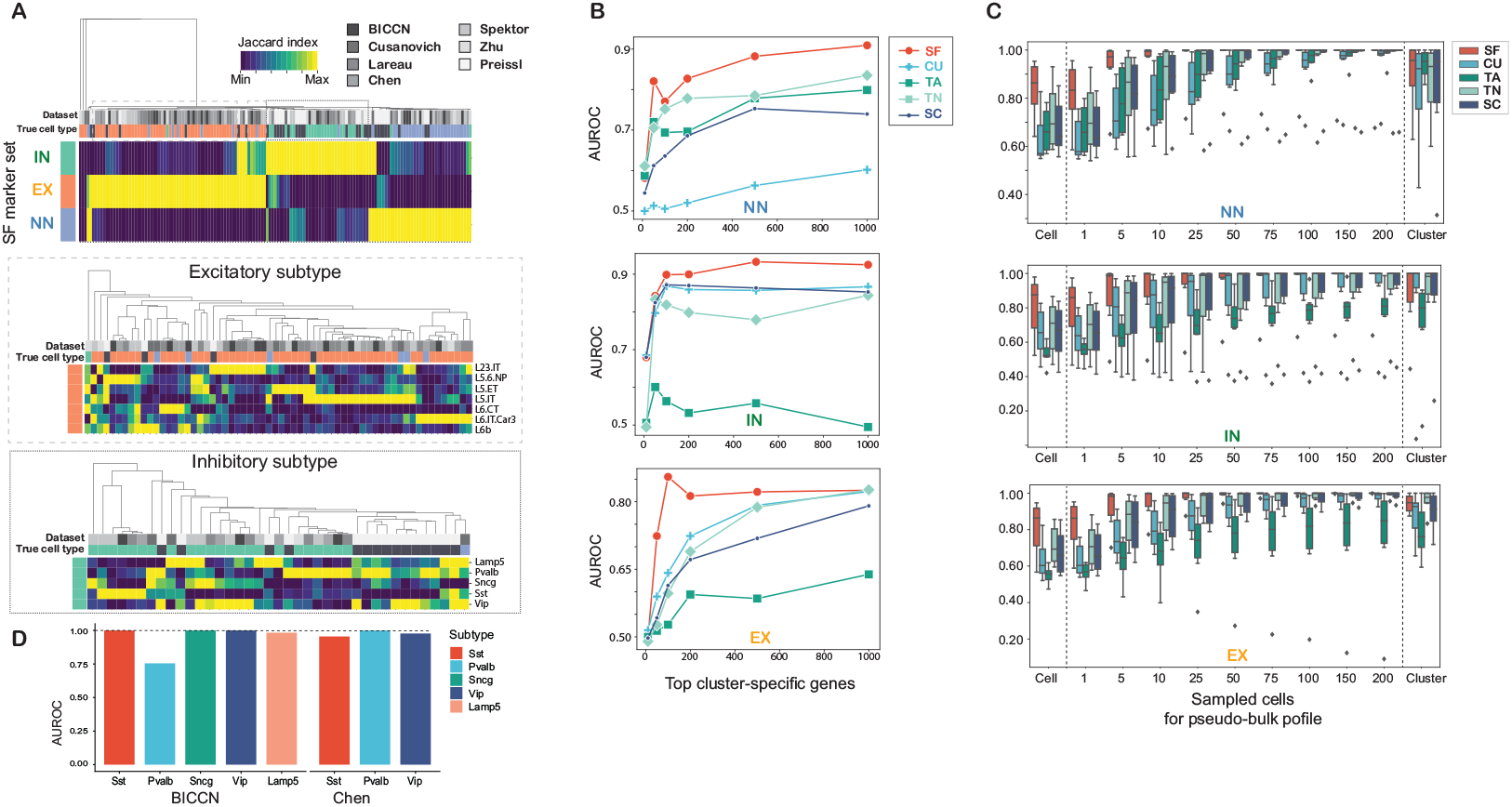
Performance of cluster-level annotation based on qualitative and quantitative cell-typing strategies. (A) The heatmaps of normalized Jaccard indices for meta-analytic marker sets and all clusters from seven scATAC-seq datasets including one dataset without cell labels (shown in dark gray). Each panel shows the scores for the SF sets of major three cell types (top), subtypes of excitatory neurons (middle) and inhibitory neurons (bottom). (B) AUROCs of major cell-type classification for six scATAC-seq datasets at cluster-level. For each cluster, a top group of cluster-specific genes is selected based on Wilcoxon’s rank sum test within each dataset (shown in x-axis). Then the Jaccard indices normalized among the three cell types are used to compute AUROCs for NN, IN, and EX cell types. (C) AUROCs of NN, IN, and EX cell-type classification within each dataset using an aggregated marker gene activity. Each boxplot represents the raw prediction accuracy of cell-level and cluster-level annotation as well as down-sampling simulation of pseudo-bulk profiles from randomly sampled cells (also see the Method section). (D) AUROCs of inhibitory neuronal subtype classification for the BICCN and Chen datasets using SF marker sets for the corresponding subtypes.

While cell-type annotation using a cluster-specific gene list is a simple and effective approach, it relies on clusters partitioning the cells appropriately into types and may be sensitive to over- or under-clustering. For that reason, we evaluated another cell-type approach by computing the absolute (rather than relative) strength of marker gene activities, which are also called module scores. This generates a single cell-type score for each cell group that can then, again, be summarized by performance as AUROCs (Fig. 3C). For cell-level annotations, we computed AUROCs from individual cell profiles, while for cluster-level annotations we used the average pseudo-bulk profiles of the cells that belong to the same cluster. Although there are only small differences in AUROCs for the cluster-level classification, the SF marker set outperformed at individual cell-level classification with a median AUROC around 0.85. Moreover, to show the robustness of marker set performance without a class imbalance problem, we constructed simulation data of 100 average profiles for each major cell type over a specific number of cells randomly sampled from the original datasets. Along with the increase of the AUROCs with the number of cells, the SF marker set is found to show the most stable prediction accuracy for cell-type classification. In summary, we found that our meta-analytic marker gene set could improve the robustness of major cell-type classification at a variety of cell resolution.

### 2.5 The robustness and feasibility of subtype classification can be improved using meta-analytic marker gene sets

In addition to the three major cell types, we can iteratively use the SF marker sets to perform rare cell- or subtype classification as shown in Figure 3A. We first extracted two groups with the higher Jaccard scores for either of IN or EX marker sets, then computed the number of overlapping genes again between each SF subtype marker set and cluster-specific genes. The cell-typing performance of several inhibitory subtypes (Sst, Pvalb, Sncg, Vip, and Lamp5) was validated for the BICCN and *Chen* datasets, which both contain the clusters associated with those subtypes (Fig. 3D). In the BICCN dataset, the AUROCs of five inhibitory neuronal subtypes at the smallest cluster level are 1.0, 0.755, 1.0, 1.0, and 0.983, respectively. These clusters, except for Pvalb, are considered to be substantially distinctive within the BICCN dataset using SF subtype marker sets. In the *Chen* dataset, the AUROCs of Sst, Pvalb, and Vip of inhibitory subtypes are all higher than 0.955. Since the SF marker sets were constructed independently from the scATAC-seq datasets, the use of meta-analytic marker sets is a promising approach to enable robust cell typing even for neuronal subtypes at the cluster-level.

Furthermore, by carefully examining the consistency of the signals in Figure 3A, some clusters from other datasets show enrichment for multiple marker sets. This suggests the heterogeneity of those clusters and our cell typing at individual cell level would be able to detect the existence of mixed subtype populations. Finally, the clusters of the *Preissl* dataset, whose “true” labels are not available in this study, also show an exclusive signal enrichment for the SF major cell- and subtype marker sets. This, too, indicates their applicability to labeling unknown clusters.

### 2.6 Comprehensive assessment of cell-type classification and marker sets with machine learning classifiers

We next ask whether more sophisticated prediction methods, such as supervised learning, can further improve performance with our marker genes. To determine the degree to which robust markers facilitate more sophisticated cell typing, we performed a comprehensive assessment of scATAC-seq cell-type classification at the individual cell level. Because of the extreme heterogeneity of the scATAC-seq datasets, it is rarely possible to select the most robust approach for not only clustering but also cell-type characterization. Instead of exploring just the best combination, we aim to address the question of whether the suitable feature selection based on marker genes is still critical beyond the differences in the test datasets, training datasets, or prediction methods.

We applied a variety of supervised learning methods for scATAC-seq, such as raw signal aggregation (as used in the previous section), machine learning (ML) classifiers, and joint clustering methods. Importantly, raw signal aggregation of the marker set is the only method that is applicable without a reference training dataset. This provides scope for methods with more parameter optimization to improve performance, although the parameter optimization can sometimes reduce robustness.

As ML classifiers, we applied four different classifiers (e.g., Logistic regression, support vector machine(SVM)) and trained them using the BICCN scRNA-seq data (*RNA atlas*) or other scATAC-seq datasets (*Consensus*). As joint clustering methods developed for the integration of scRNA-seq, BBKNN [22] and Seurat [23] were selected. Further details on optimization and evaluation are described in the Methods section (also see Supplementary Fig. 2).

Figure 4A shows the summary of the AUROCs for NN cell-type classification at the individual cell level. The prediction performance highly depends on the dataset quality or similarity as shown in a clear contrast between the rows: when the dataset is too deviated from others, no single method or training condition seemed to work at a practical level. In Figure 4B, the top 10 combinations in terms of average AUROCs are extracted. Most of the combinations are based on ML optimized on the Consensus (using all other datasets for training) although also included is one combination of the raw signal aggregation and SF marker sets. The two best methods are the Logistic regression classifier trained on all genes, and Alternate Lasso trained on SF marker genes. With respect to the marker sets selected, SF and TN gene sets are dominant within the top 10 combinations while only one combination utilizes all gene sets. This suggests the utility and stability of redundant marker gene sets as a kind of feature selection method against the dataset-dependent variability.

**Fig.4.**
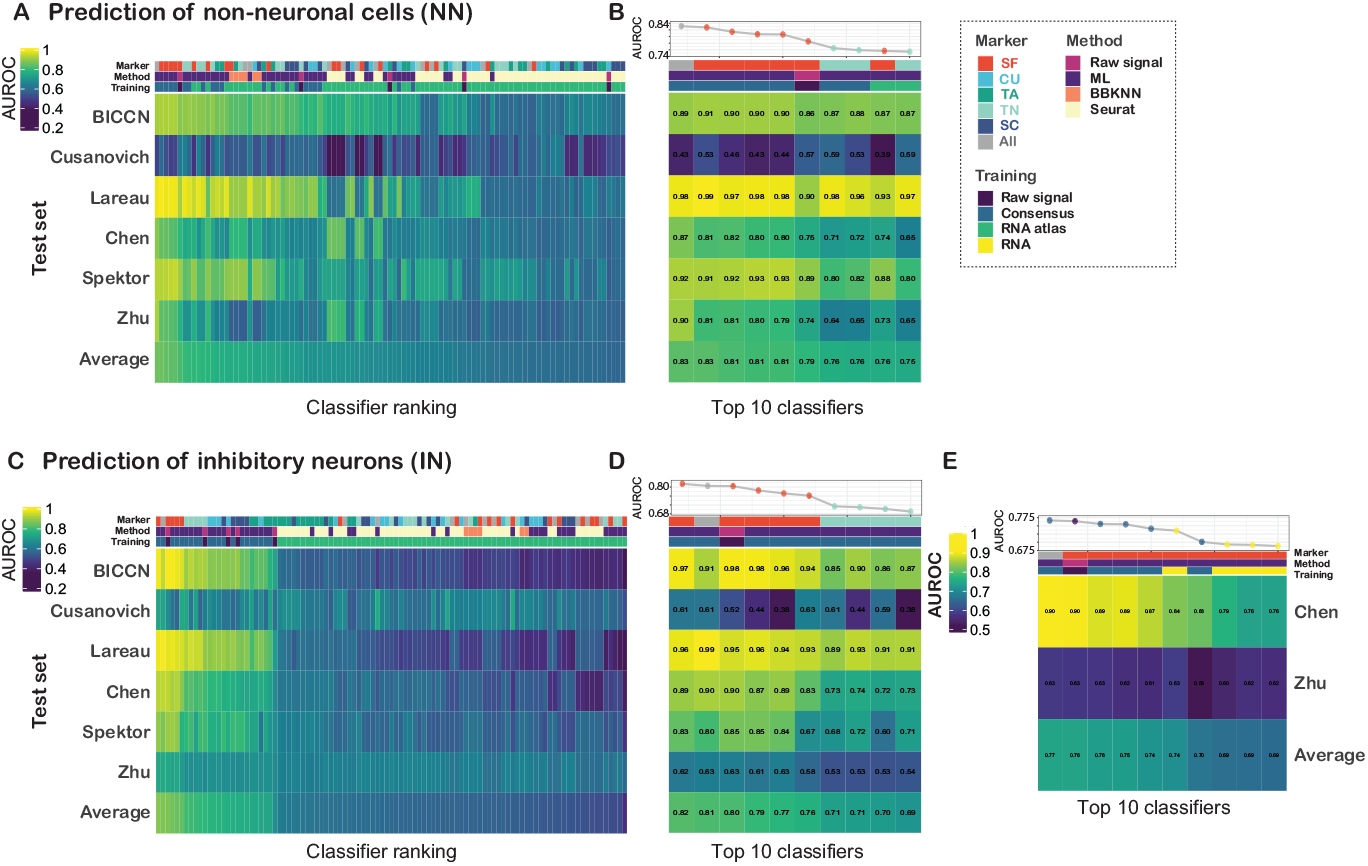
Comprehensive comparison of cell-type classification at individual cell level using a combination of marker genes and supervised learning methods. (A) and (B) AUROCs of six datasets and those average for NN cell-type classification at each cell level for all demonstrated combinations (A) and top 10 combinations according to the average AUROCs (B). The columns shown at right represent the marker set, supervised learning method, and training dataset used to construct each classifier. (C) and (D) AUROCs of 6 datasets and those average for IN cell-type classification at each cell level for all demonstrated combinations (C) and top 10 combinations according to the average AUROCs (D). (E) AUROCs of two joint profiling datasets and those average for IN cell-type classification at each cell level for top 10 combinations according to the average AUROCs. This result contains the combinations using available scRNA-seq data of each dataset as a training data.

Next, the prediction performances for IN cell-type classification are visualized (Fig. 4C and D). Unlike the result of NN cell-type classification, the AUROCs of the top and bottom combinations are clearly distinct. Specifically, the combinations based on Consensus training or raw signal aggregation show apparently higher AUROCs than those using the RNA atlas from the BICCN dataset. As previously, the top 10 combinations exploit the combinations involving the SF and TN marker sets as well as only one set using all genes. Indeed, even the simple raw signal aggregation method from the SF marker gene sets is ranked in the top 10 regardless of the target cell types while that with other marker sets are not.

Additionally, we examined the classification performance using a transcriptome-based reference from the exact same sample (named *RNA* training) to show the potential of the scRNA-seq reference. Figure 4E shows the top 10 combinations that performed best for two joint profiling datasets. The top 10 ranks are occupied by methods of ML classifiers optimized by Consensus and RNA reference data, in addition to raw signal aggregation for the SF marker gene set.

In summary, our comprehensive assessment strongly suggests that consensus training using other scATAC-seq data and simple aggregation of large marker sets are comparably powerful for major cell-type classification. Although optimization based on the independent reference scRNA-seq was less powerful, training on the joint-profiled scRNA-seq shows a comparable prediction performance with consensus training. More importantly, in all cases, the choice of marker genes most strongly characterized the performance of a method/data/feature combination, suggesting the wide-applicability of robust marker gene sets for integrative analyses and interpretation of the resultant cell-type specific ATAC-seq profiles for regulatory inference, as described next.

### 2.7 Meta scATAC-seq server with deep learning prediction enhances cell-type specific motif analysis

For future integrative analysis, we have published a new Meta scATAC-seq server at https://gillisweb.cshl.edu/Meta_scATAC/ to make all collected data and marker genes in this study available. In this server, the average read count of each genomic location can be visualized in a genome browser for certain cell types or datasets. The accessibility signals can be aggregated for not only the cluster-derived cell types provided by the authors (which are not unified across the studies) but also the top 500 cells of each subtype by calculating the marker activity of SF.

Furthermore, to assess the potential of genomic sequences to regulate the cell-type specific cis-regulatory programs, we integrated predicted chromatin accessibility data using a sequence-based deep CNN trained on the BICCN scATAC-seq data. Specifically, we generated a multitask classification dataset that consists of cell-type specific pseudo-bulk chromatin profiles. The pseudo-bulk profiles were then used to generate a dataset of 5kb DNA with a corresponding label vector that specifies whether the DNA is accessible or not in the BICCN dataset (see Methods). We constructed a custom CNN with a Basset-like architecture [24] and trained it to take DNA as input and simultaneously predict chromatin accessibility across each cell-type (Fig. 5A). We found that the CNN’s classification performance on test data (i.e. all data from held-out chromosomes: 1, 3, and 5) had good predictive power with an area under the precision-recall curve (AUPR) of 0.539 on average across each celltype – this is a large improvement upon DeepSea’s AUPR of 0.444 across 125 chromatin accessibility datasets [25], most of which derive from cell lines.

**Fig.5.**
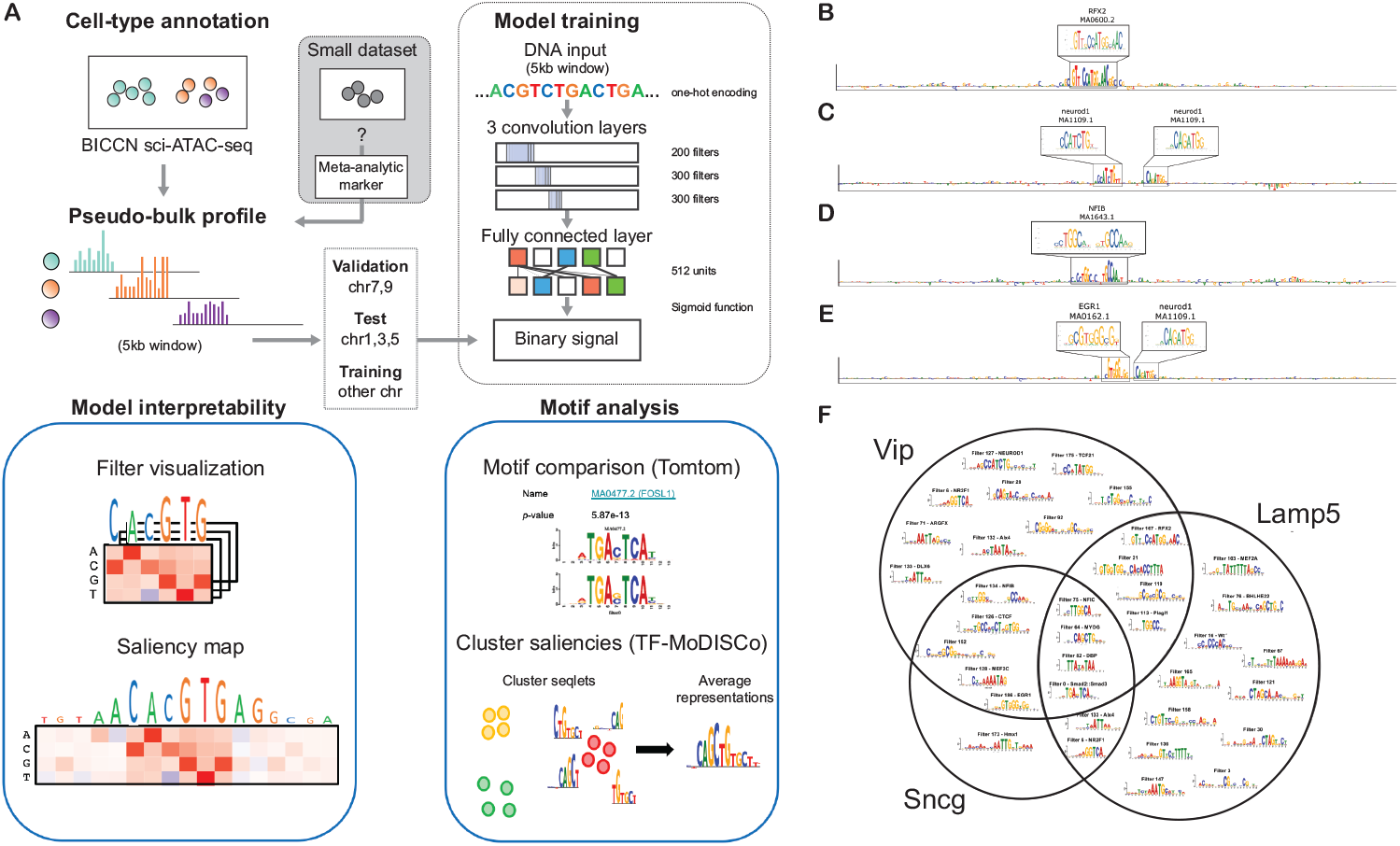
(A) Deep learning analysis workflow. The CNN model was trained to take DNA sequence as input and predict the sequence’s accessibility in each cell type. Model interpretability is accomplished by visualizing layer convolutional filters and by cell-type specific saliency analysis for a given sequence. Each can be used to visualize motif features extracted by the model. Motif analysis is then performed by employing two ways of motif search: a motif comparison search against a database of known motifs (i.e. JASPAR) using Tomtom or via TF-MoDISCo, which splits saliency maps, clusters them, and provides an averaged representation. Motif analysis of BICCN scATAC-seq pseudo-profiles (B-F). Sequence logos of saliency maps from a CNN model trained at the sub-cell type level: (B) Lamp5, (C) Vip, (D) Sncg, and (E) Vip. Only 210 positions out of 5kb is shown for visual clarity. The known motifs from the JASPAR database are annotated with a box above each saliency plot, labelled with a putative motif name and JASPAR ID. (F) Venn diagram of motifs enriched in each cell type. The filter representations are shown, while the cell type-specific motif enrichment was determined with TF-MoDISCo and the motif annotations were given by statistically significant matches to the JASPAR database using Tomtom.

For model interpretability, we performed filter visualization and attribution methods, both of which are common techniques in genomics [26]. We compared filter representations against the 2020 JASPAR vertebrates database [27] using Tomtom [28]. We found that 36% of the filters match known motifs, which is seemingly a low number considering that when applying a similar network to the Basset dataset, our CNN yields a higher match fraction of about 62% [29]. Using an attribution method called saliency analysis [30], which takes the gradient of a given class prediction with respect to the inputs, we can generate sequence logos of the importance of each nucleotide in a given sequence (see Methods). Within the 5kb binned sequences, we often find that small patches within the attribution maps highlight known motifs either alone (Fig. 5B), in combinations with their reverse complements (Fig. 5C-D), and with other partners (Fig. 5E).

To demonstrate the efficiency of meta-analytic marker genes for the interpretation of a deep CNN model, we examine enriched patterns within the attribution maps, specifically nearby meta-analytic marker gene sets, using a clustering tool for attribution maps TF-MoDISCo [31]. Figure 5F highlights a Venn diagram of the motifs enriched in different cell types: Lamp5, Vip, and Sncg. Many motif representations were shared between all three cell types, including the binding motifs of NFIC, MYOG, DBP, and FOSL1. Vip and Lamp5 had many unique motifs enriched near meta-analytic marker genes, while Sncg only had a single enriched motif identified by TF-MoDISCo. This is consistent with the strong overlaps among these cell-types within the observed transcriptional hierarchy, where there is mixing across types when defined purely by expression clusters [32]. A full list of the motif matches for each filter is provided as Supplementary data 7. We also found that many TF-MoDISCo cluster representations, which were also supported by convolutional filter representations, do not have any correspondence to a known motif in the JASPAR database – these were labeled by just their filter name. This was expected to an extent as the JASPAR database is not complete and the ability to analyze cell type-specific regulatory regions within the brain emerged recently with the advent of scATAC-seq data.

## 3 Discussion

In this study, we examined the usability of meta-analytic marker genes for scATAC-seq cell-typing at a variety of levels of granularity. We found that a robust marker gene set optimized for multiple scRNA-seq data produced high performance at resolutions from cell to cluster level. Interestingly, feature selection via marker gene sets substantially improved neuronal and non-neuronal cell-type prediction even without sophisticated supervised learning methods. The choice of marker gene sets was a major driving factor of the prediction performance rather than that of classification strategies. We also demonstrated the potential power of this strategy for regulatory inference from scATAC-seq experiments once heterogeneous data has been partitioned into cell-type specific pseudo-bulk profiles. The resultant “cleaned” profiles can be used to train a CNN model to learn a relationship between input DNA sequence and its accessibility for a given cell-type – we showed this with the BICCN dataset. Interpreting the trained CNN revealed learned motifs that were enriched near the marker gene sets in a subtype-specific manner, suggesting the existence of cell-type specific regulation for meta-analytic marker genes in the motor cortex. The straightforward feasibility of using robust marker gene sets for accurate subtype cell-typing within scATAC-seq data opens up many important downstream possibilities, most clearly condition- and subtype-specific regulatory network discovery, as demonstrated in our own deep learning analysis.

ScRNA-seq has proven to be a remarkably effective technique for the characterization of cell-types within the brain, shedding new light on decades old questions regarding the form, function, and organization of cell-types [33]. In turn, the complexity of the brain has made it one of the strongest use cases for single cell technologies. While cell-typing and characterization have been major success stories, understanding the regulatory basis of the observed cell-types remains an important challenge [9]. One of our important contributions is to demonstrate just how effective marker gene selection can be for brain epigenetic data. The cell-type characterization based on marker genes has long been a mainstay of wet-lab biology but typically focused on specificity, rather than comprehensiveness [34]. In contrast, high-throughput single-cell methods have generally preferred methods that rely on information distributed across a large fraction of genes. Our analysis suggests that a middle ground of picking redundant marker sets meta-analytically satisfies a number of important constraints: high classification performance, generalization, simplicity and straightforward interpretability. While we see dramatic differences from dataset to dataset, feature selection appears to be the critical determinant for accurate cell-typing, as opposed to more complicated modeling of the way those features interact (which is less likely to generalize). Because marker sets can be derived from high-performing scRNA-seq data, we exploit all the existing success of cell-typing efforts there to inform the interpretation of scATAC-seq data. Importantly, the utility of feature selection for consistent annotation is likely to remain even as wet-lab technology improvements (such as paired scRNA-seq and scATAC-seq) will make clustering cells within a given dataset less challenging. The significance of marker set selection is also highlighted by the improved interpretability it offered when we turned to model the cell-type specific regulatory programs through deep learning.

To predict chromatin accessibility across different cell-types from just the DNA sequence, CNNs have demonstrated a remarkable ability [24, 25, 35]. ScATAC-seq provides an opportunity to study cell-type specific regulatory programs in heterogeneous tissues, such as the immune cells [36] and the brain (this study), using CNNs. By “purifying” scATAC-seq data by robust cell typing, the accessibility signal is expected to become more reliable. This may explain why our CNN yields improved performance both in classification accuracy and interpretable motif representations. Since many accessible sites are shared across different cell-types, these “overlapping” regions may not necessarily contain the information we desire, that is to know which motifs drive cell-type specific regulation. Hence, it remains a challenge to decipher which motifs are relevant for cell-type specificity. Our approach was to explore the enrichment of motifs nearby meta-analytic marker genes that are celltype specific, which shifts the distribution of transcription factors that are learned genome-wide to the ones that regulate genes of a given cell-type. Indeed this approach reveals many known motifs (and some putative novel ones). Moving forward, it would be beneficial to follow up this work to try to decipher which proteins bind to these motifs in each cell-type and explore which non-marker genes they regulate.

## 4 Methods and Materials

### Meta-analytic marker genes for mouse brain

For robust and accurate cell-typing of single-cell data, we developed a method, MetaMarkers, and used it to define a new meta-analytic marker gene set, called *SF* marker set [15], derived from multiple scRNA-seq datasets from the BICCN. The SF marker set is the expanded marker set that includes co-expressed genes of marker genes. Additionally, *SC* is a subset of SF but restricted to have the same number of genes as that of CU to assess the importance of the number of genes, rather than the protocol by which the marker genes were obtained. The overlap of each marker gene is shown in Figure 2D and F (also see Supplementary Fig. 1).

### Mouse brain scATAC-seq datasets

From the BICCN collection, we obtained single-cell combinatorial indexing ATAC-seq (sci-ATAC-seq) data which consists of 4 batches with a transcriptome reference of SMART-seq v4 scRNA-seq data (the cell number after filtering is 6,278) for the mouse primary motor cortex region [16]. Both datasets are available from the BICCN data portal https://biccn.org. Moreover,we collected scATAC-seq datasets of the mouse brain published on Gene Expression Omnibus (GEO). Specifically, read count matrices and metadata of 6 scATAC-seq studies were downloaded from GEO. The corresponding GEO IDs of the collected studies areGSE100033 [1], GSE111586 [12], GSE123576 [37], GSE127257 [38], GSE126074 [17], and GSE130399 [18]. From the Paired-seq datasets of GSE130399, the one for an adult mouse cerebral cortex sample is applied in this study. To convert read counts to gene activities, we used the gene structure information from an Ensembel GTF file for GRCm38 as of Nov. 2018. Each genomic feature in the original study was then assigned to the closest TSS found in the GTF file. A gene activity estimation was carried out by summing the read counts of all assigned features within the 10kb upstream or downstream from the TSS of all transcripts of the same gene id. For the datasets whose feature is peak-based, the locations of each peak center were used to associate each feature and gene. A general pre-processing, filtering, clustering, and detection of cluster-specific genes was performed on a SCANPY platform [39].

### Assessment of cell-typing for scATAC-seq

We performed a comprehensive assessment of cell-typing for six well-annotated scATAC-seq datasets using a different combination of supervised learning methods, training sets, and marker gene sets. A graphical outline is shown in Supplementary Figure 2.

#### Supervised learning methods

The methods used in this study are classified into three categories: *raw signal, ML classifiers*, and *joint clustering methods*. Raw signal methods predict each cell type based on the raw signal scores computed by aggregating the read counts for the genes included in each biomarker gene set. This method is the only method that does not require any training dataset except for a marker gene set. ML classifier methods consist of four popular ML classifiers applicable to supervised learning of scATAC-seq cell-typing. Specifically, SVM, random forest, logistic regression with L1 regularization (Logistic regression), and a variant of logistic LASSO “Alternate Logistic LASSO” [40] are included in this category, which is expected to be more robust for sparse data. Due to the limitation of computational resources, only Logistic regression was carried out for the prediction using all genes and the other three classifiers were applied with the feature selection based on the marker gene set. The last category is a joint clustering method, in which a test and training dataset is reanalyzed independently, then jointly clustered two datasets to associate each cell in the test set with the annotated cells in the training dataset. We chose Seurat [23] and BBKNN [22] for a comparison referring to the results of the previous study of an integrated analysis of single-cell atlases [14]. To compute AUROCs from the results of BBKNN, we implemented our own script to compute the scores for a cell-type prediction by counting the nearest-neighbor cells for each cell type.

#### Training set

The training set is applied in four different ways. Raw signal methods use gene activity profiles from the test scATAC-seq dataset only for selected biomarker genes. Consensus methods use the scATAC-seq datasets except for the one used as a test set. For this prediction, the prediction scores are computed by the classifiers trained on each training set. Those scores are averaged to compute the final prediction scores after normalization within each dataset. RNA atlas methods use the BICCN scRNA-seq data as a training set to optimize the parameters or infer the nearest-neighbor cells. For the datasets based on joint profiling methods, we also carried out ML-based supervised learning using scRNA-seq from the same dataset, named “RNA” training.

#### Gene set selection

In addition to the five marker gene sets collected from the previous studies, we performed the supervised learning with all detected genes if an optimization process is feasible. Specifically, supervised learning based on all genes was demonstrated for Logistic regression from ML classifiers and both joint clustering methods.

### Supervised learning by machine learning classifiers

To carry out supervised learning, we implemented a workflow of optimization of ML classifiers using the scikit-learn library. The parameters used for each classifier are as follows: degree is 3 and kernel is set to an rbf kernel for SVM, n_estimators is set to 100 for RF, and C is set to 1.0 (default) for Logistic regression. Other parameters are set to the default values. For Alternate Lasso, we implemented an original classification function in which the best and alternate predictors are averaged with different weights. In this study, we extracted top *n* predictors at maximum and summed their predictions with the weight 1/*n*, where n is set to 5.

### Joint clustering methods for an integrative analysis of single-cell omics datasets

BBKNN was applied to the pair of scATAC-seq and the BICCN scRNA-seq dataset after applying a general normalization for the scRNA-seq dataset by a Scanpy function “normalize_per_cell”. The parameter for k-nearest neighbor used in the BBKNN algorithm was set to *k* = 5, 10, 20, 30 with and without a graph trimming option. We also run Seurat v3.2.2 to align the same dataset combinations as used for BBKNN. The alignment of two datasets was done via FindTransferAnchors and TransferData functions using canonical correlation analysis by setting a parameter reduction=‘cca’.

### Sequence analysis of accessible sites with deep learning

#### Data for binary classification

Given the 5kb bins, each of which contains a single bulk accessibility profile value, we binarized each label with a threshold of 0.02, above which is given a positive label, and below is given a negative label. We then filtered sequences with no positive labels across all classes. Sequences were converted to a one-hot representation with 4 channels (one for each nucleotide: A, C, G, T) and a corresponding label vector with either a 0 for negative labels or 1 for positive labels. We split the data into a validation set (chromosomes 7 and 9; *N* = 24, 908), test set (chromosomes 1, 3, and 5; *N* = 14, 670), and training set (all other chromosomes; *N* = 274, 689).

#### Model

Our CNN model takes as input a 1-dimensional one-hot-encoded sequence with 4 channels, then processes the sequence with three convolutional layers, a fully-connected hidden layer, and a fully-connected output layer that have sigmoid activations for binary predictions. Each convolutional layer consists of a 1D cross-correlation operation followed by batch normalization [41], and a non-linear activation function. The first layer used an exponential activation, which was previously found to encourage first layer filters to learn interpretable motif representations and also improves the overall interpretability with attribution methods [29]; while the rest used a rectified linear unit. The first convolutional layer employs 200 filters each with a size of 19 and a stride of 1. The second convolutional layer employs 300 filters each with a size of 9 and a stride of 1. And the third convolutional layer employs 300 filters each with a size of 7 and a stride of 1. All convolutional layers incorporate zero-padding to achieve the same output length as the inputs. The first two convolutional layers are followed by max-pooling layer of window size 10, and the last one is followed by a global average pooling layer. The fully-connected hidden layer employs 512 units with rectified linear unit activations. Dropout [42], a common regularization technique for neural networks, is applied during training after each convolutional layer, with a dropout probability set to 0.2 for convolutional layers and 0.5 for fully-connected hidden layers.

#### Training

All models were trained with mini-batch stochastic gradient descent (mini-batch of 100 sequences) with Adam updates [43] with a decaying learning rate using a binary cross-entropy loss function. The initial learning rate was set to 0.001 and decayed by a factor of 0.3 if the model performance on a validation set (as measured by the Pearson correlation) did not improve for 7 epochs. Training was stopped when the model performance on the validation set does not improve for 25 epochs. Optimal parameters were selected by the epoch which yields the highest Pearson correlation on the validation set. The parameters of each model were initialized according to Glorot initialization [44].

#### Filter visualization

To visualize first layer filters, we scanned each filter across every sequence in the test set. Sequences whose maximum activation was less than a cutoff of 50% of the maximum possible activation achievable for that filter in the test set were removed [24, 45]. A subsequence the size of the filter centered about the max activation for each remaining sequence and assembled into an alignment. Subsequences that are shorter than the filter size due to their max activation being too close to the ends of the sequence were also discarded. A position frequency matrix was then created from the alignment and converted to a sequence logo using Logomaker [46].

#### Saliency analysis

To test the interpretability of trained models, we generate saliency maps [30] by computing the gradients of the predictions with respect to the inputs. Saliency maps were multiplied by the query sequence (times inputs) and visualized as a sequence logo using Logomaker [46].

## Acknowledgements

We thank Dr. Rongxin Feng, Dr. Song Chen, and Dr. Chenxu Zhu for providing us with a script and metadata for their studies used in our metaanalysis. We also would like to thank Dr. Sukalp Muzumdar, Jonathan Werner, John Hover, and other lab members for their thoughtful comments.

## Author contributions

JG conceived the project. JG and RKK designed computational experiments. RKK performed computational experiments. PK, ZT, CR, and RT developed and interpreted deep learning models. RKK wrote the initial draft of the manuscript with assistance from JG, SF and PK. SF generated meta-analytic marker lists. All the authors read and approved the final manuscript.

## Data availability

Source codes and marker gene sets are available at https://github.com/carushi/Catactor Pseudo-bulk profiles of all collected data are published at Meta scATAC-seq server https://gillisweb.cshl.edu/Meta_scATAC/.

## Author Biographies

**Risa Karakida Kawaguchi** is a postdoc fellow at Cold Spring Harbor Laboratory, US. Her research interests include an integrative analysis of multi-omics data and epigenetic regulation of gene expression. **Ziqi Tang** is a Ph.D student at Cold Spring Harbor Laboratory, US. Her research interests include the application and interpretation of deep learning models for epigenetics data. **Stephan Fischer** is a postdoc fellow at Cold Spring Harbor Laboratory, US. His research interests include statistical properties and co-expression network of scRNA-seq datasets. **Chandana Rajesh** is a Ph.D student at Stony Brook University, US. Her research interests include application of deep learing for genomics and neuroscience data. **Rohit Tripathy** is a postdoc fellow at Cold Spring Harbor Laboratory, US. His research interests include the development of new machine learning tools and application of deep generative models to extract novel insights from genomic data. **Peter K. Koo** is an assistant professor at Simons Center for Quantitative Biology, Cold Spring Harbor Laboratory, US. His research interests include deep learning-based sequence analyses and their interpretability improvement. **Jesse Gillis** is an associate professor at Department of Physiology and Donnelly Centre for Cellular & Biomolecular Research Department, University of Toronto, Canada. He is also an adjunct professor at Cold Spring Harbor Laboratory, US. His research interests include network analysis for neuropsychiatric data and co-expression network within scRNA-seq.

