## Supplementary data for "Learning single-cell chromatin accessibility profiles using meta-analytic marker genes"

Risa Karakida Kawaguchi[1], Ziqi Tang[1], Stephan Fischer[1], Chandana Rajesh[1], Rohit Tripathy[1], Peter K. Koo[1], and Jesse Gillis[1,2]*
1 Cold Spring Harbor Laboratory, 1 Bungtown Rd, Cold Spring Harbor, New York, 11724, USA.
2 Department of Physiology and Donnelly Centre for Cellular & Biomolecular Research Department, University of Toronto, Ontario, M5S 3E1, Canada.

### Supplementary Figure 1

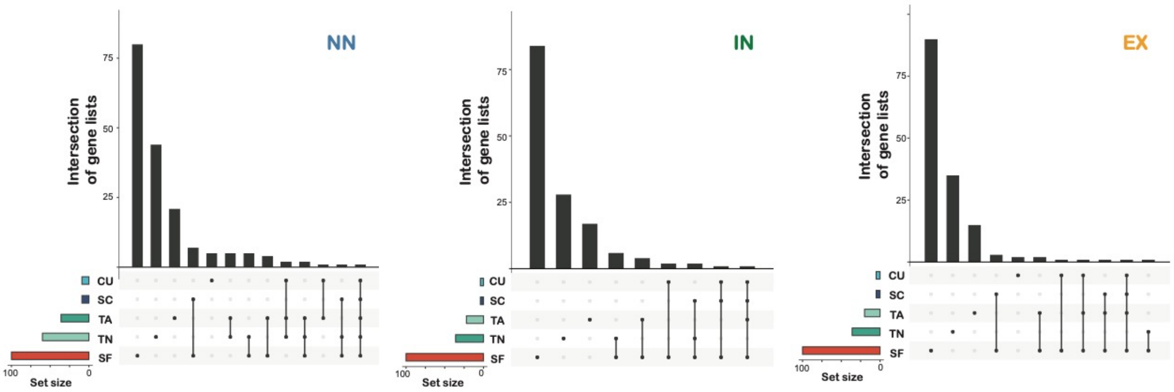


(A) The AUROC distribution of major cell-type classification for the BICCN scATAC-seq dataset using the gene activity of each single gene. The AUROCs lower than 0.5 are converted into the opposite direction so that all AUROCs are no less than 0.5 by the formula $\max(1.0 - \mathrm{AUROC}, \mathrm{AUROC})$. (B), (C), and (D) The top 1,000 features for cell-type classification of the BICCN dataset based on different features or different problems. (B) Cell-typing for each cell using a single gene activity. (C) Cell-typing for each cluster using a single gene activity from the averaged profile. (D) Cell-typing for each cell using an accessibility of each 5kb genomic bin at cell-level annotation. The inset plot shows the $\log_{10}$ p-value after Bonferroni multiple correction computed by Fisher's exact test from the binarized accessibility for each genomic bin.

### Supplementary Figure 2

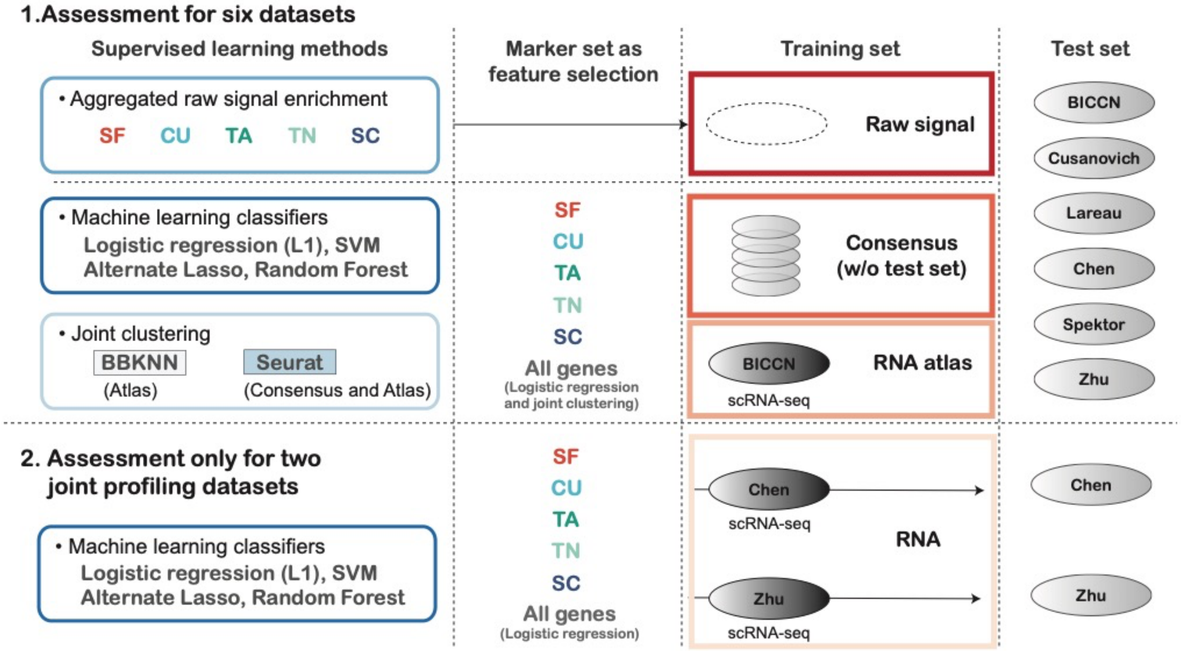


A workflow of a comprehensive assessment of scATAC-seq cell-type classification at an individual cell level.

### Supplementary Figure 3

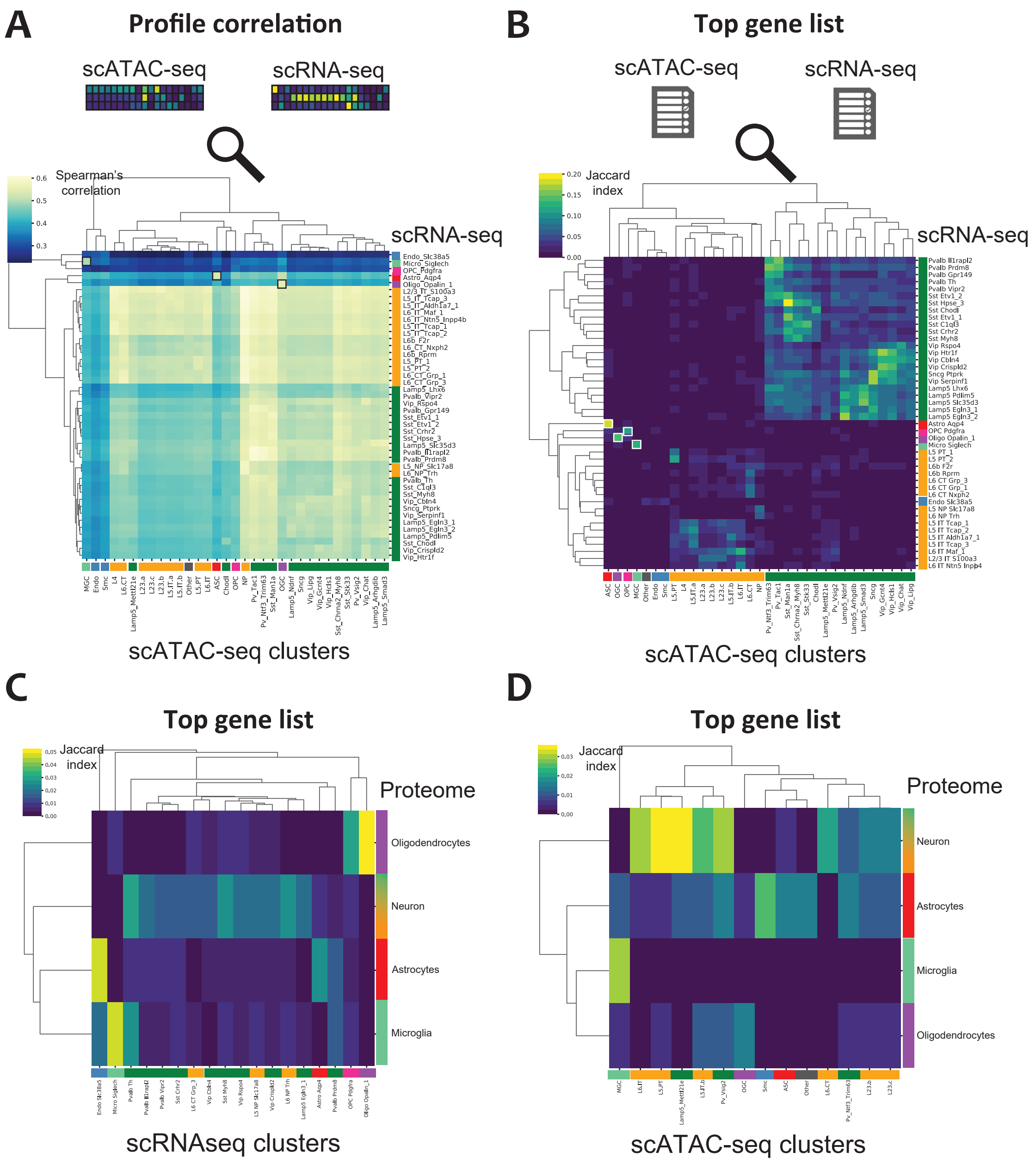


(A) The heatmaps of Pearson's correlation coefficient between the average gene expression and gene activity of whole genes from scRNA-seq and scATAC-seq clusters, respectively. (B) Normalized Jaccard indices for the top 100 cluster-specific genes for each cluster between the BICCN scRNA-seq and scATAC-seq datasets. (C) and (D) The heatmaps of normalized Jaccard indices between the top cluster-specific genes from the proteome dataset (Sharma K., et al. 2015) and scRNA-seq (C) and scATAC-seq (D).

### Supplementary Figure 4

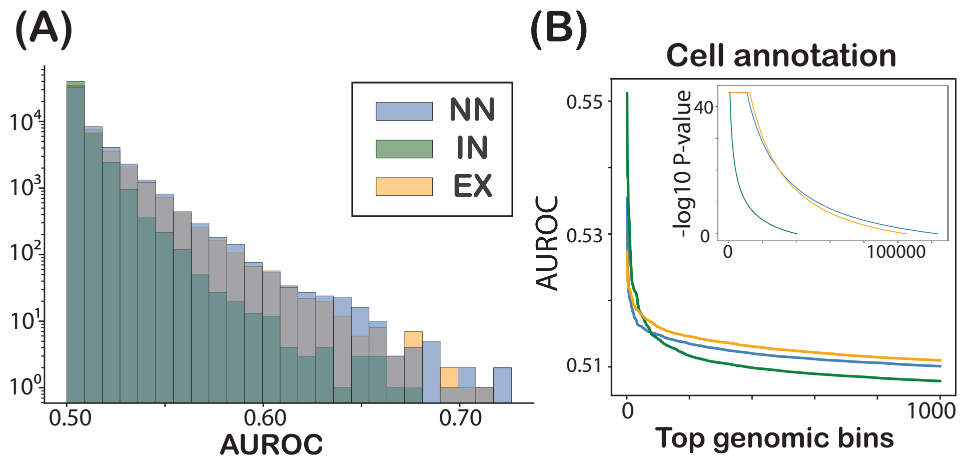


(A) The AUROC distribution of major cell-type classification for the BICCN scATAC-seq dataset using the gene activity of each single gene. The AUROCs lower than 0.5 are converted into the opposite direction so that all AUROCs are no less than 0.5 by the formula $\max(1.0 - \mathrm{AUROC}, \mathrm{AUROC})$. (B), The top 1,000 features for cell-type classification of the BICCN dataset based on different features or different problems. Cell-typing for each cell using an accessibility of each 5kb genomic bin at cell-level annotation. The inset plot shows the $\log_{10}$ p-value after Bonferroni multiple correction computed by Fisher's exact test from the binarized accessibility for each genomic bin.

### Supplementary Figure 5

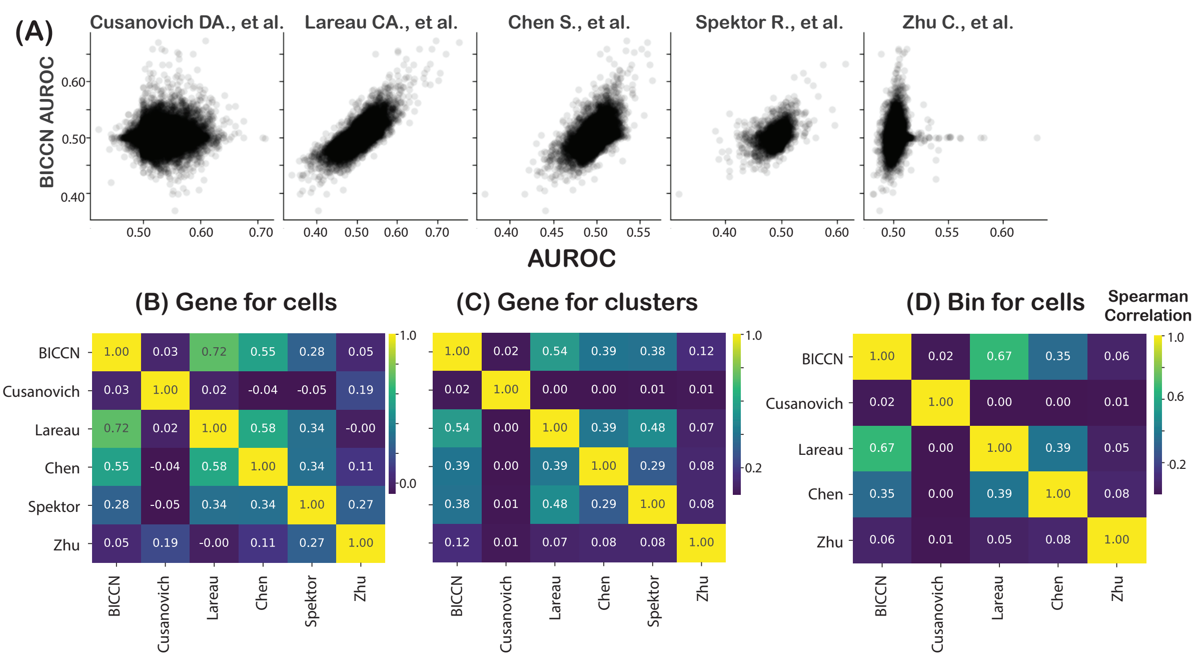


Consistency of cell-type classification by each single feature across the datasets.
(A) Scatter plots of the AUROCs of IN cell-type classification within the BICCN dataset (y-axis) and other 5 scATAC-seq datasets (x-axis). (B), (C), and (D) Spearman's correlation coefficients of the AUROCs based on each gene activity for cell-level annotation (B), averaged gene activity for each cluster (C), and signal activity for each 100kb genomic bin (D).

### Supplementary Figure 6

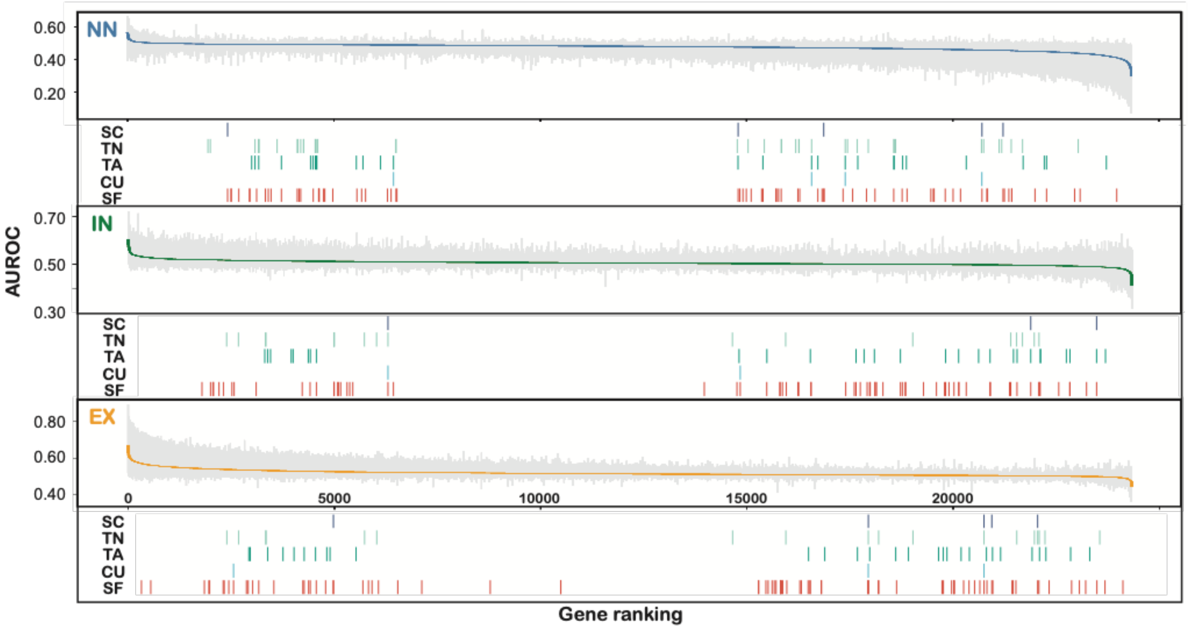


The distribution of mean, min, and max AUROCs for three cell-type classification using the estimated gene activity across the six scATAC-seq datasets with the information of marker gene annotation. The solid line shows the average of the AUROCs for the six datasets while the top and bottom of gray lines correspond to the maximum and minimum of AUROCs for each gene. The top, middle, and bottom panel indicate the AUROCs for NN, IN, and EX cell-type classification. In the bottom rectangle of each panel, short vertical bars are shown at the location of genes listed in the marker set for each cell type.

### Supplementary Figure 7

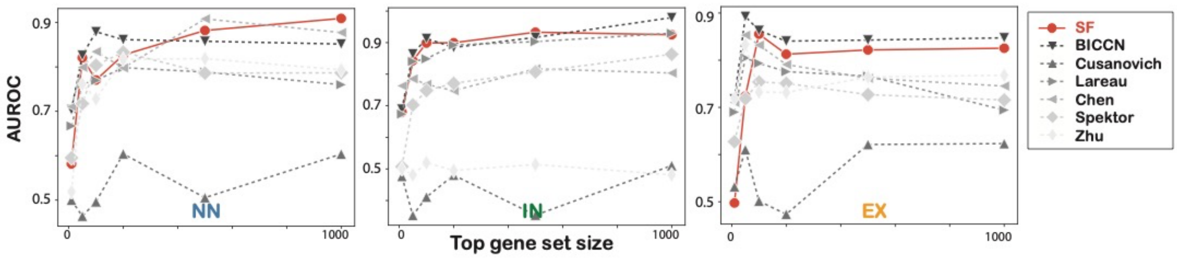


AUROCs of cluster-level cell-type classification by computing normalized Jaccard scores for top cluster-specific genes and marker gene sets. As a marker set, the AUROCs of top 1,000 cell-type specific genes derived from each dataset for non-overlapping datasets are compared with those of the SF marker gene sets across all well-annotated datasets.
